# A tetracycline-inducible promoter for gene expression in Gram-negative bacteria including *Acinetobacter baumannii*

**DOI:** 10.64898/2026.10.09.758105

**Authors:** Nicholas A. Stella, Ronit K. Ginde, Raj K. Ginde, Rachel C. Calvario, Yohei Doi, Robert M. Q. Shanks

## Abstract

Carbohydrate inducible promoters on expression plasmids are widely used for biotechnology and basic science research. However, using carbohydrates for inducers has drawbacks including influencing bacterial metabolism, the inducer being used as a carbon source by the bacteria, high background expression, and only a subset of cells experiencing promoter activation. Furthermore, some bacteria including the pathogen *Acinetobacter baumannii* currently lack good carbohydrate-inducible promoters. To address this issue, a tetracycline-inducible promoter, *P_tetB_*, isolated from *A. baumannii* was cloned and evaluated in a number of important bacteria: *Acinetobacter baumannii*, *Escherichia coli*, *Klebsiella pneumoniae*, *Pseudomonas aeruginosa*, *Pseudomonas fluorescens*, and *Serratia marcescens*. A highly sensitive *luxCDABE* reporter was used to evaluate promoter activity. *S. marcescens* strains with elevated and reduced cyclic-AMP levels were used to evaluate the impact of cAMP on P*tetB* function. The *P_tetB_* promoter was induced over 800-fold in *A. baumannii* and 10-750-fold inducible in the other tested species. Background expression was lower from *P_tetB_* in the pseudomonads than previously described carbohydrate inducible promoters. Data indicated that the P*tetB* promoter system may be a useful tool for future basic science research with the tested bacteria.

## 1. INTRODUCTION

Carbohydrate-inducible promoters are widely used in the biological sciences. However, there are certain drawbacks such as the organism consuming the inducer as a carbon source. The impact of cyclic-AMP (cAMP) on these promoters can also be a challenge when studying biological systems in which cAMP levels are altered between groups. For some organisms, commonly used carbohydrate-inducible promoters fail to function or have very high levels of background in the absence of inducer. For example, despite the importance of *Acinetobacter baumannii* as an opportunistic infection, there are few studies describing a useful inducible system. Commonly used carbohydrate-inducible promoters were not inducible (1). Exceptions were reported by Lucidi, et al 2019 (2), who developed plasmids with a DNA-damage inducible *uvrA* promoter, ethanol-induced promoters *adhP* and *yahK*, and an iron limitation-inducible promoter *basA*. Though these may be useful under certain circumstances, there are experimental conditions where use of UV, mitomycin C, iron chelation, and possibly ethanol could be deleterious through impacts on bacterial genetic fidelity and growth. Furthermore, these promoters were approximately 2 to 4-fold inducible at the highest inducer levels (2), which is relatively modest. Therefore, we sought a promoter system that had a high level of inducibility and low background levels in some commonly studied Gram-negative bacteria including *A. baumannii*.

In this study, we evaluated a tetracycline-inducible promoter with accompanying transcriptional repressor sourced from *A. baumannii*. This promoter was cloned on broad host-range vectors and tested in a variety of opportunistic pathogens using a luminescent reporter.

## 2. MATERIALS AND METHODS

### 2.1. Strain and culture conditions

Bacterial strains are listed in Table 1. Cultures were propagated in lysogeny broth (LB) (3) at 30°C or 37°C with aeration using a floor shaker or tissue-culture rotor. Gentamicin was used for plasmid maintenance at 10-30 µg/ml depending on the host strain. Plasmids were introduced into *E. coli* EC100D (Lucigen) by electroporation (4), whereas transfer to the remaining species was accomplished by conjugation. *E. coli* strain HB101 (5) served as a helper strain for triparental matings. Following conjugation, donor and helper strains were counterselected using ampicillin (150 µg/ml), chloramphenicol (30 µg/ml), nalidixic acid (20 µg/ml), or tetracycline (10 µg/ml).

**Table 1.** Strains used in this study.

| Strain | Description | Source |
| --- | --- | --- |
| ATCC 17978 | *A. baumannii* | (24) |
| B5 | *A. baumannii* multi-drug resistant clinical isolate | (1) |
| S17-1 $\lambda$ *pir* | *Escherichia coli* | (25) |
| Kp2<br>(ATCC43816) | *Klebsiella pneumoniae* | (26) |
| Pf0-1 | *Pseudomonas fluorescens* | (27) |
| UCBPP-PA14<br>(PA14) | *P. aeruginosa* | (28) |
| Db11 | *Serratia marcescens* | (29) |
| K904 | *Serratia marcescens* | (30) |
| PIC3611 | *Serratia marcescens* | (31) |
| Δ*crp* | *Serratia marcescens* with mutation of *crp* | (13) |
| Δ*cpdS* | *Serratia marcescens* with mutation of *cpdS* | (14) |

### 2.2. Relative light unit (RLU) determination

RLU were determined as previously described (1). Bacterial colonies were used to inoculate 4.5 ml of LB broth with 0.5 ml of inducer (as noted in the figure legend) or water as a negative control. After growth at the indicated time and temperature, 150 µl aliquots were transferred to 96 well plates with black sides and optical polystyrene bottoms (Nunc) and read with Spectramax M3 and L plate readers to assess optical density (OD_600_) and luminescence respectively. RLU was defined as luminescence signal divided by optical density.

### 2.3. Molecular biology

Plasmids were generated using yeast gap repair, in which PCR amplicons or synthetic DNA fragment were co-transformed with a linearized shuttle vector into *Saccharomyces cerevisiae* strain INVSc1 (Invitrogen) (6, 7). Table 2 lists the plasmids used and or generated in this study. Plasmids were validated by PCR and full plasmid sequencing (PlasmidSaurus).

**Table 2.** Tetracycline-inducible plasmids made for this study.

| Plasmid no. | Relevant feature <sup>a</sup> | Replicon |
| --- | --- | --- |
| pMQ609 | $P_{tetB} - gfp$ | *oriRK2, ColE1* |
| pMQ707 | $P_{tetB} - gfp$ | *p15a, oripW1266* |
| pMQ790 | $P_{tetB} - luxCDABE$ | *pBBR1* |
| pMQ808 | $P_{tetB} - MCS^b$ | *p15a, oripW1266* |
| pMQ909 | $P_{tetB} - MCS^b$ | *pBBR1* |

<sup>a</sup> all have a gentamicin resistance marker, *aacC-1*
<sup>b</sup> MCS, multicloning site included BamH1, HindIII, SpeI, and SphI sites

To make pMQ609, a tetracycline inducible promoter and tetracycline repressor gene originally from *A. baumannii tetB* were cloned from pBCSK(L-R)*tetR* (8) with primers 3989 and 3990. This was moved into pMQ123i (6) cut with BamH1 and PmlI in order to replace P*_BAD_*. To make pMQ707, the *tetB* promoter (P*_tetB_*) was amplified with primers 919 and 4025, and used to replace P*_BAD_* in pMQ560 that had been cut with BspE1 and Sal1. To make pMQ790, the *rhaB* promoter on pMQ688 was replaced with P*_tetB_* from pMQ707. The *tetB* promoter was amplified with primers 5524 and 5525 and the amplicon was recombined with pMQ688 that had been digested using EcoRI and PvuI. To make pMQ808 and pMQ909 the *luxCDABE* reporter was replaced by a multicloning site amplified from pMQ782 (1) or pMQ790 with primers 923 and 5635 or with the synthetic DNA fragment 5597 (Table 3). The pMQ707, pMQ808, and pMQ909 plasmid sequences were submitted to GenBank (ID: PP526038.1, PP737597, and QB173169).

**Table 3.**
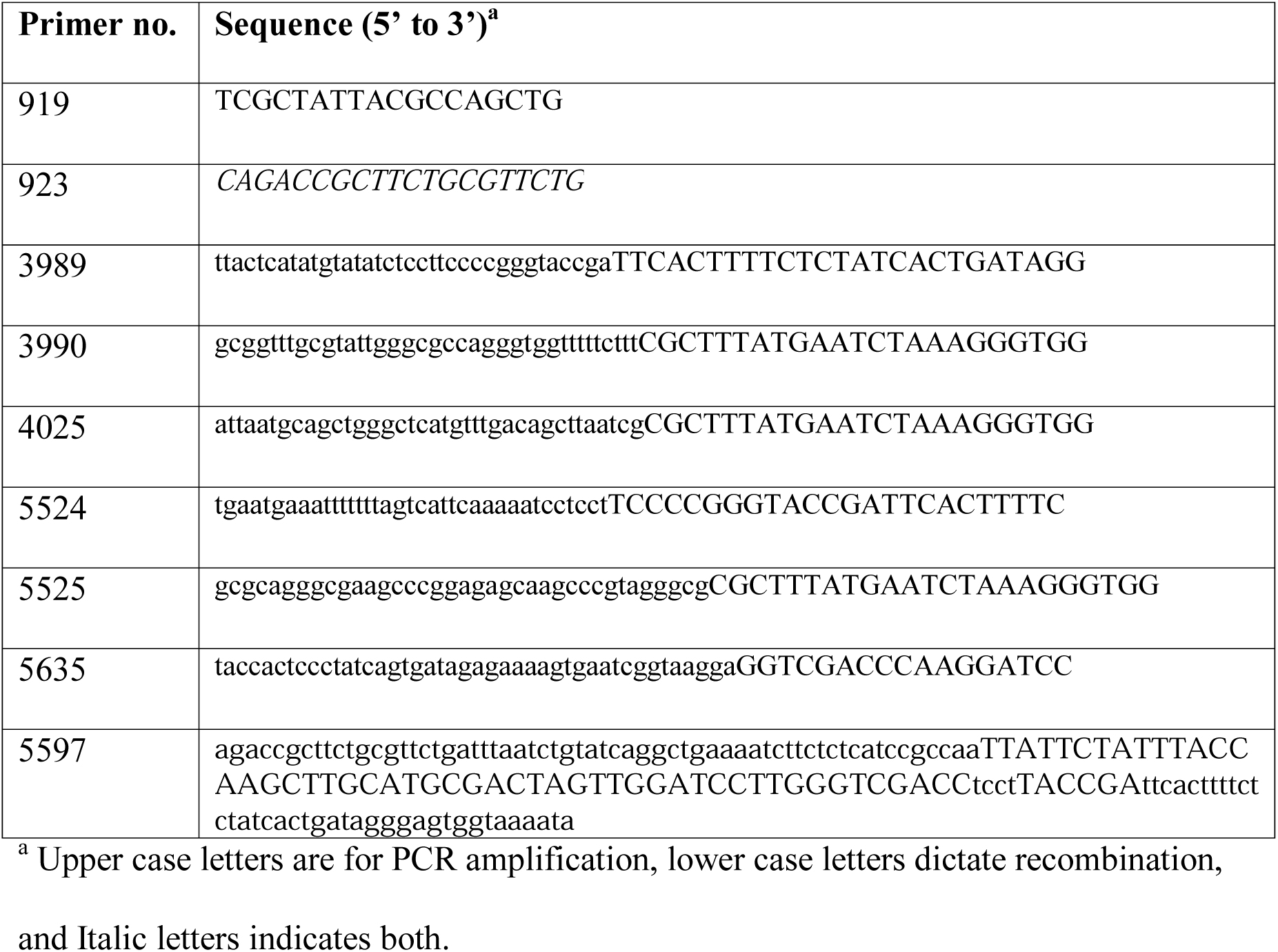
Oligonucleotide primers used in this study.

### 2.4. Statistical analysis

Statistical analysis was performed using GraphPad Prism software. One-way ANOVA with Tukey’s multiple-comparison test or Student’s T-test were used as indicated in the figure legends.

## 3. RESULTS

### 3.1. Testing a tetracycline inducible promoter from *A. baumannii*

In a previous study of carbohydrate-inducible promoters in a variety of Gram-negative opportunistic pathogens, none were inducible in *A. baumannii* (1). In a search for an inducible promoter system, we evaluated a tetracycline inducible promoter originally from *A. baumannii* as a candidate inducible promoter (8). This included a *tetR* regulator gene and predicted promoter with two operator sequences in a region spanning 813 bp (Figure 1A). The *tetR* gene codes for a protein that is 100% identical at the amino acid level to previously discovered tetracycline resistance regulators (9), and this protein has been used in a bistable circuit in *E. coli* indicating that it will function in species other than *A. baumannii* (10).

**Figure 1.**
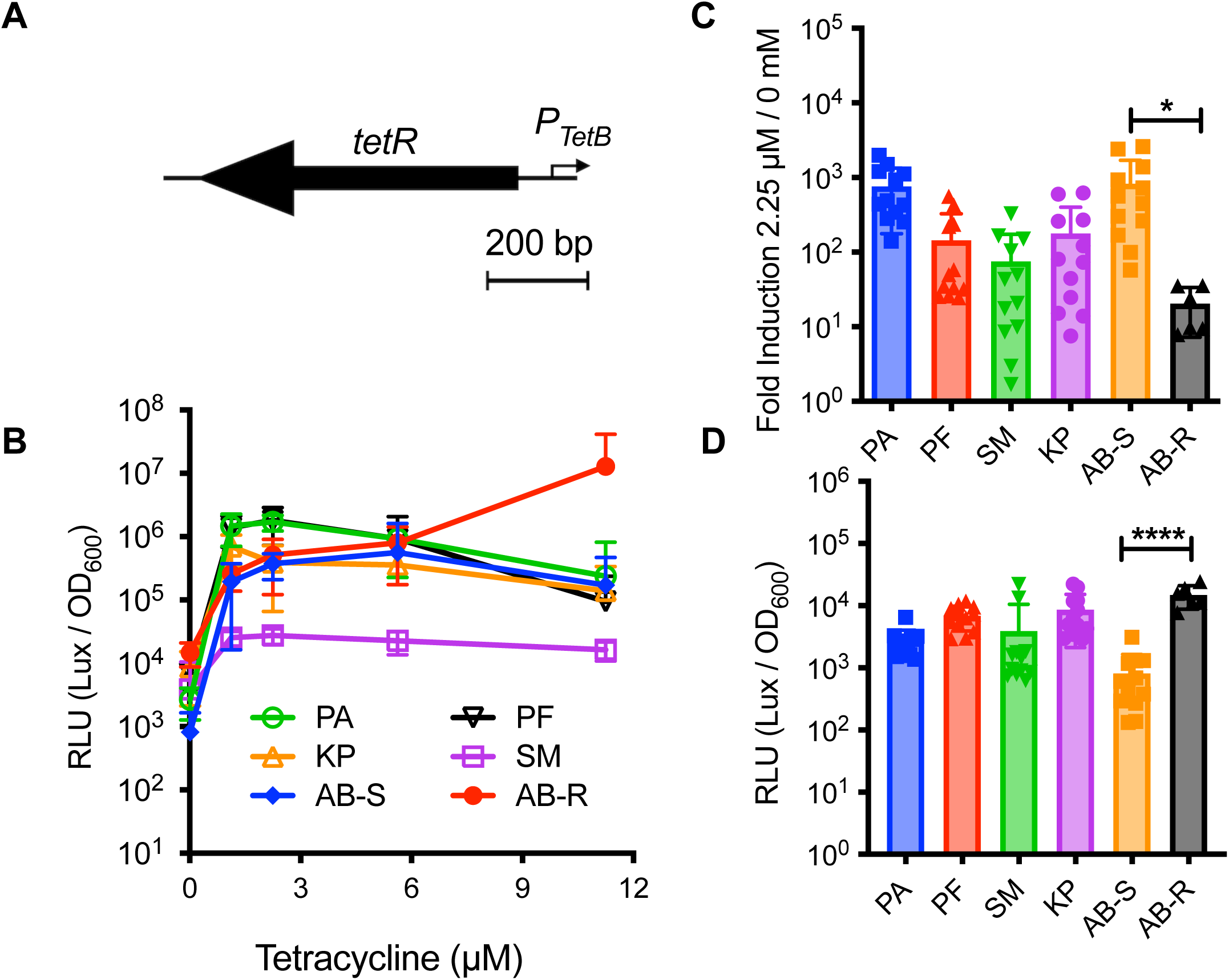
A tetracycline inducible promoter was inducible in *A. baumannii* and other bacteria. **A.** The regulatory region from a tetracycline resistance marker used to drive *luxCDABE* in this study. **B.** Luminescence from P*_tetB_* in a variety of bacterial species over a concentration gradient of tetracycline (n=6-12). **C.** Fold induction of each strain growth with tetracycline at 2.25µM compared to no inducer. **D.** Background luminescence in the absence of tetracycline. Mean and SD are shown. PA, *P. aeruginosa* PA14; PF, *P. fluorescens*, Pf01; SM, *S. marcescens* Db11; KP, *K. pneumoniae* Kp2; AB-S, *A. baumannii* ATCC17978, and AB-R, *A. baumannii* B5. Asterisks show significant differences determined by ANOVA with Tukey’s post-test. *, p<0.05; ****, p<0.0001. Bacteria were grown at 30°C. LUX indicates luminescence; OD_600_ culture turbidity.

The *P_tetB_* promoter was evaluated using a *luxCDABE* reporter with plasmid pMQ790 in *A. baumannii* and the other bacteria of interest over a range of concentrations (Figure 1B). In this set of experiments, the inducer was added at time zero and measured growth and luminescence following overnight growth. The fold induction was generally highest at 1.125-2.25µM of tetracycline for each strain tested (Figure 1B). For *A. baumannii* strain ATCC 17978 that is tetracycline susceptible (AB-S) and *P. aeruginosa* (PA14), over 700-fold induction was measured (Figure 1C). This was followed by *P. fluorescens* and *K. pneumoniae* with over 100-fold induction, and *S. marcescens* (Db11) with approximately 75-fold induction (Figure 1C).

To test the applicability of this promoter to other strains of *A. baumannii*, a multidrug resistant clinical isolate B5 (AB-R) was used to evaluate the efficacy of *P_tetB_* promoter in a tetracycline-resistant isolate (Figure 1B). The fold increase was 20±13 which was much lower than for the susceptible ATCC strain suggesting that tetracycline resistance status may influence the *P_tetB_* promoter strength (Figure 1C).

Background luminescence from the *P_tetB_* promoter was determined and found to be reasonably low (Figure 1D). Notably, it was >10-fold lower for tested *Pseudomonas* species compared to the carbohydrate-inducible promoters in our prior study including arabinose and rhamnose inducible promoters under identical conditions (1).

### 3.2. Evaluation of P*_tetB_* over time at 37°C

The prior experiments evaluated various promoters at a single time point after overnight growth at 30°C. To expand on this, experiments were performed to evaluate earlier time with a subset of organisms when grown at 37°C. Bacteria were grown overnight without induction and subcultured with inducer at 1.125 µM for tetracycline added at time 0 h. Luminescence and optical density were read over time 5h and the next day (Figure 2A-C).

**Figure 2.**
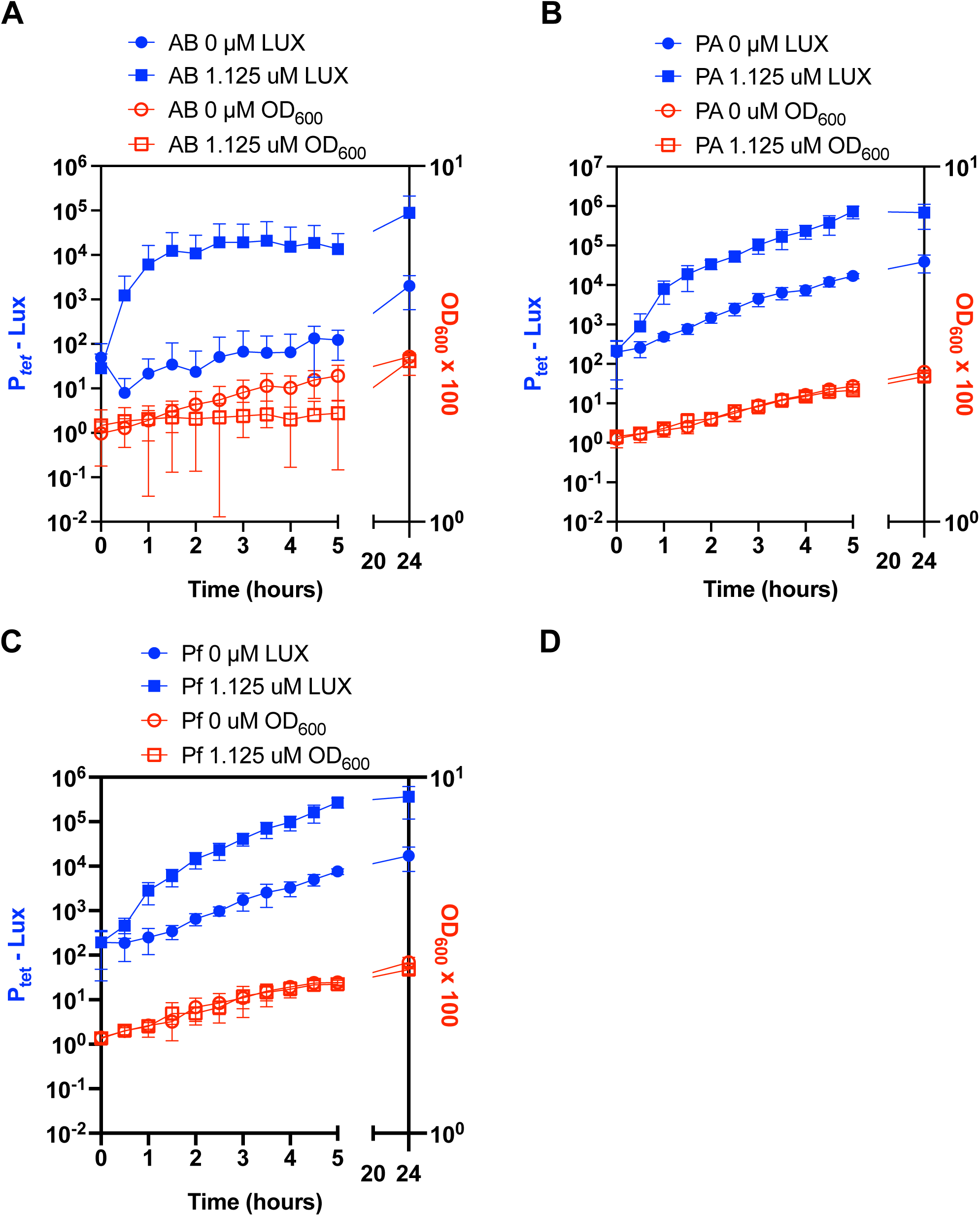
Induction of the P*_tetB_* promoter over time at 37°C. **A-C.** Growth and P*_tet_*-dependent luminescence of bacteria over time. AB, *A. baumannii* strain ATCC 17978; PA, *P. aeruginosa* strain PA14; Pf, *P. fluorescens* strain Pf0-1. Inducer and vehicle negative control were added at time = 0. Mean and SD are shown, n=4-6. LUX indicates luminescence; OD_600_ culture turbidity. The concentration of tetracycline is listed in the legend.

Luminescence from the P*_tetB_* promoter was measured in *A. baumannii* strain ATCC17978 and it increased 43-fold by 0.5h and more than 725-fold by 3.5h compared to time 0h (Figure 2A). Without inducer, there was little induction compared to time 0h over the first 5h. Growth, however, was reduced with 1.125 compared to 0 µM tetracycline, and further concentration optimization is likely important in a strain-by-strain manner.

Early induction of the P*_tetB_* promoter was also evaluated in two *Pseudomonas* species (Figure 2B-C). In both *P. aeruginosa* and *P. fluorescens* induction was observed by 30 minutes in the tetracycline-induced cultures. At 5h, luminescence was 44 and 36-fold higher for PA14 and Pf0-1 in the tetracycline-induced group than negative control group, and 233 and 92-fold higher at 5h compared to 0h in the tetracycline group. Tetracycline did not impact culture optical density of these pseudomonads when used at 1.125 µM.

### 3.3. Use of anhydrotetracycline as an inducer

We tested anhydrotetracycline as an alternative inducer as it may have less antibacterial effect (11). Figure 3 shows the ability of 0.1 µM of anhydrotetracycline (ATC) for growth and induced luminescence over time. ATC strongly induced P*_tetB_* in *P. aeruginosa*, *P. fluorescens*, and *A. baumannii* strains. ATC poorly induced *S. marcescens*. Both the *A. baumannii* ATCC 17978 and *S. marcescens* strains demonstrated reduced growth in the presence of ATC.

**Figure 3.**
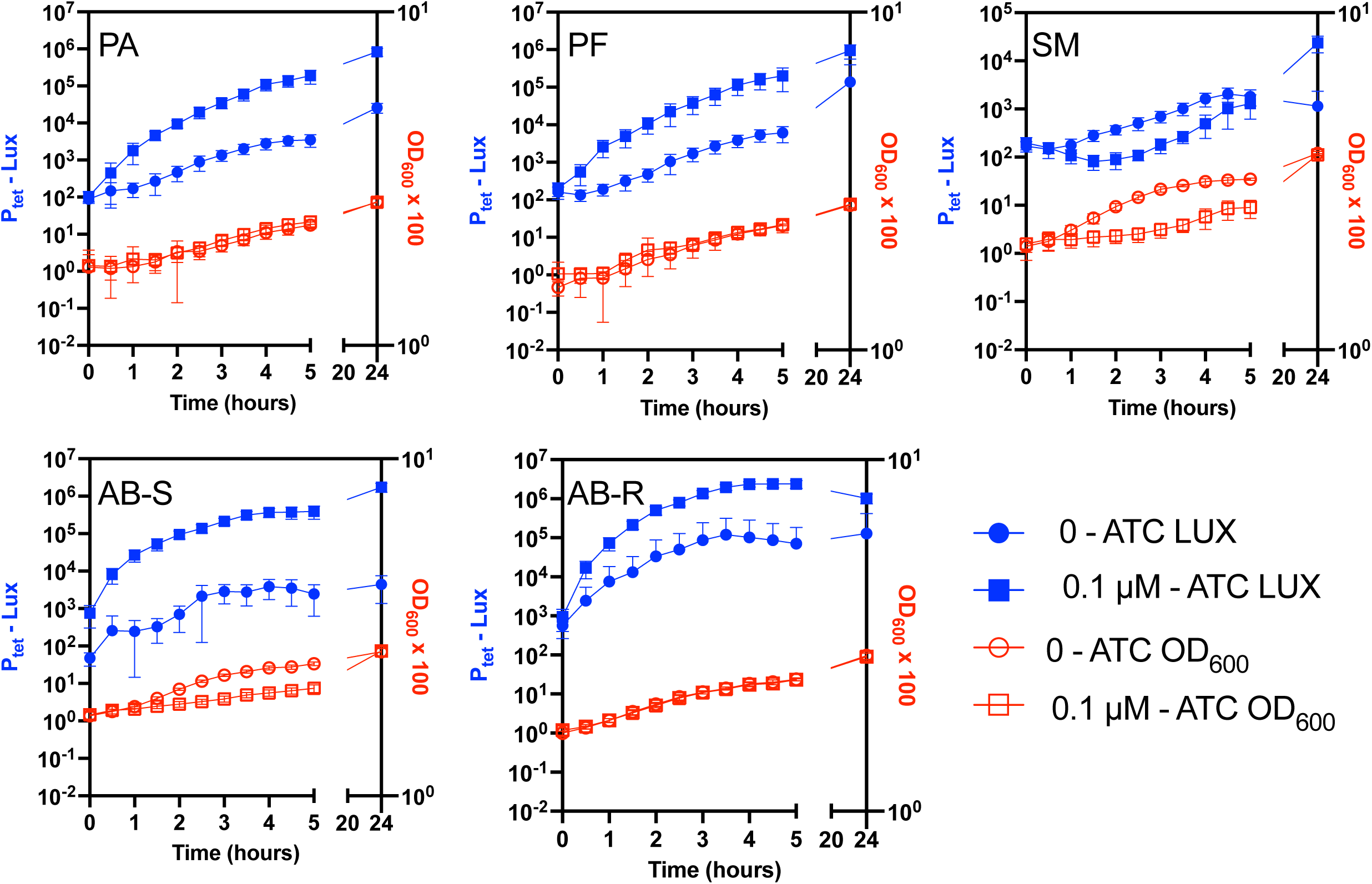
Induction of P*_tetB_* with anhydrotetracycline (ATC) at 37°C. Mean and SD P*_tet_*-dependent luminescence and growth values are shown. Inducer and vehicle negative control were added at time = 0. n=6-8. LUX indicates luminescence; OD_600_ culture turbidity. PA, *P. aeruginosa* PA14; PF, *P. fluorescens*, Pf01; SM, *S. marcescens* Db11; AB-S, *A. baumannii* ATCC17978, and AB-R, *A. baumannii* B5.

### 3.4. Evaluation of the effect of cyclic-AMP (cAMP) on P*_tetB_* expression

Some promoters, such as the *rhaB* and P*_BAD_* promoter are highly influenced by cAMP levels (12). To test the impact of cAMP on the P*_tetB_* promoter, we used isogenic *S. marcescens* strains with a mutation previously demonstrated to alter intracellular cAMP levels or response to cAMP (13, 14). The strains included the wild type PIC3611, a mutant with elevated cAMP (Δ*cpdS*), and a mutant unable to respond to cAMP, Δ*crp*. Luminescence and growth were similar among the strains when induced with tetracycline (Figure 4A-B). Though basal levels in the Δ*crp* mutant were lower than those of the other two strains. These suggest that induction of P*_tetB_* is largely cAMP-independent, at least with *S. marcescens*.

**Figure 4.**
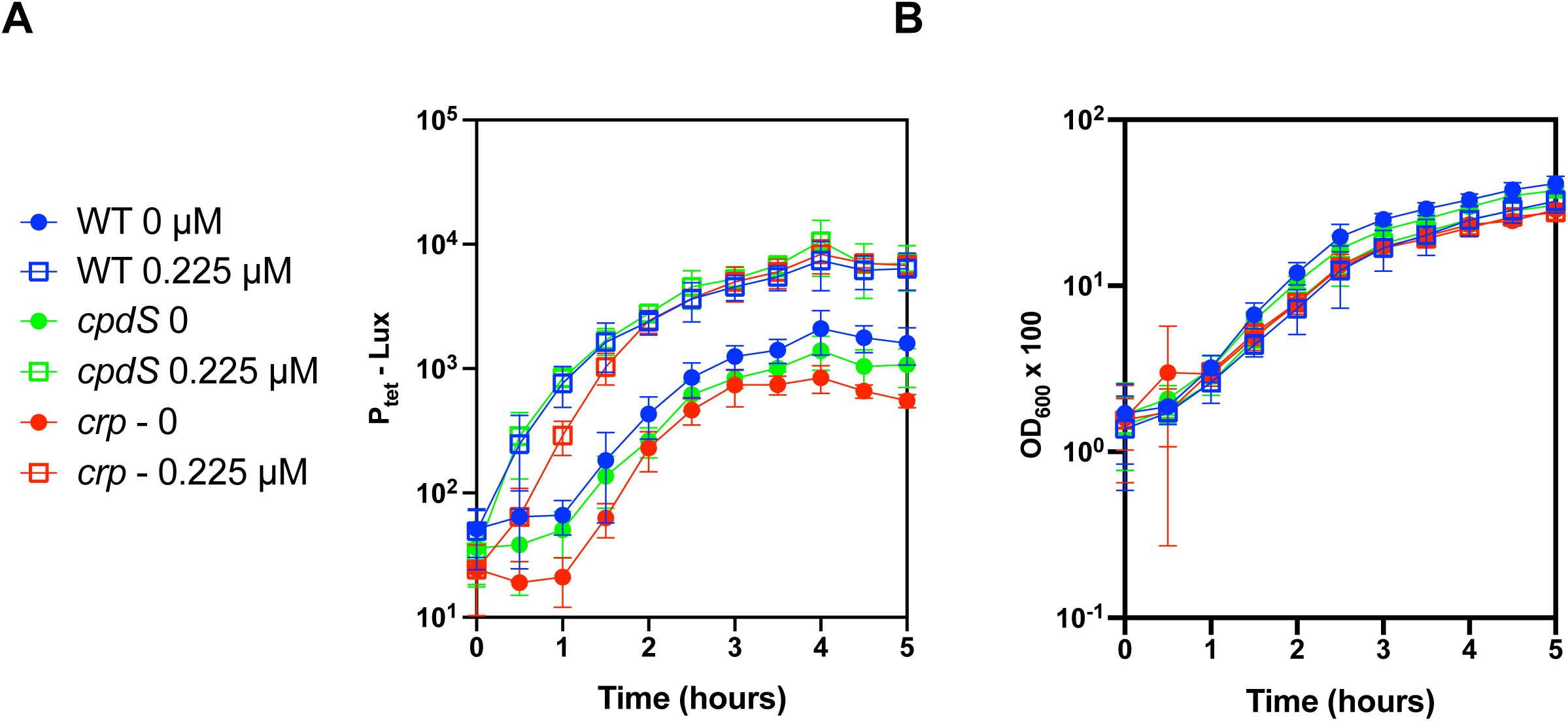
Induction of P*_tetB_* is largely cyclic-AMP independent. The concentration of tetracycline is listed in the legend. **(A)** P*_tetB_*-dependent luminescence and **(B)** growth of bacteria over time. Inducer and vehicle negative control were added at time = 0. WT, *S. marcescens* strain PIC3611; *cpdS*, PIC3611 with deletion of cAMP phosphodiesterase - elevated cAMP; *crp*, PIC3611 with deletion mutation in cAMP-receptor protein gene. Mean and SD are shown, n=4-6. LUX indicates luminescence; OD_600_ culture turbidity.

### 3.4. Generation of *P_tetB_* vectors with multicloning sites

The reporter plasmids thus far described in this study lack convenient restriction enzyme sites downstream of the tetracycline promoter. To address this, an artificial multicloning site with unique restriction sites for BamH1, HinDIII, SpeI, and SphI, a ribosome binding site, and stop codons in three frames was introduced. One was constructed in a plasmid with a p15a and pWH1266 replicon, the former for *E. coli* and related genera, and the latter for stable plasmid maintenance in *A. baumannii* (15), yielding plasmid pMQ808. A second one, pMQ909, was made with the broad host range pBBR1 replicon (16) (Table 2).

## 4. DISCUSSION

In our previous study using a variety of sugar-inducible promoters, none were functional in *A. baumannii* (1). In contrast, the data suggest that the tetB promoter was highly functional in *A. baumannii*, especially the tetracycline susceptible strain. One limitation of using tetracycline as an inducer is that it inhibits bacterial growth in susceptible strains, and appears to be less inducible in resistant bacteria. There are tetracycline variants capable of interacting with TetR with reduced antimicrobial activity, such as anhydrotetracycline that have been used with many bacteria that could mitigate this potential problem (11, 17–23). Additionally, inducer could be added after cultures reach a desired concentration for desired protein production. Another advantage of this promoter is that, at least in one bacterial species, its induction was largely cAMP-independent.

In summary, the results of this study suggest that the *tetB* promoter system isolated from *A. baumannii* should have some utility for inducible gene expression in a variety of Gram-negative bacteria. The main disadvantage is the potential growth inhibition of the inducer. However, multiple advantages include low background, potential cAMP-independence, and usefulness in *A. baumannii* and other tested Gram-negative microbes.

## 5. Acknowledgements

The funding for this study was from the National Institutes of Health grants R01EY036686 (RS), R01AI104895 (YD), and P30 EY08098 to the Department of Ophthalmology. Additional funding was given by the Eye and Ear Foundation of Pittsburgh and from an unrestricted grant from Research to Prevent Blindness, New York, NY.

